# Exploring the Impact of Apocarotenoids on Pathogenic *Fusarium oxysporum* f.sp. *lini* and Endophytic Fo47 strains

**DOI:** 10.1101/2024.11.28.625830

**Authors:** Yelyzaveta Kochneva, Marta Burgberger-Stawarz, Aleksandra Boba, Marta Preisner, Justyna Mierziak-Derecka, Anna Kulma

## Abstract

The *Fusarium oxysporum species complex* (FOSC) contains highly specific plant pathogens and some nonpathogenic strains, such as Fo47. In our work we concentrated on *Fusarium oxysporum* f.sp. *lini* (Foln), the specific flax pathogen and the endophytic strain *Fusarium oxysporum* 47 (Fo47), which is possibly protective for flax against pathogens. We investigated the influence of apocarotenoids like ionones and abscisic acid (ABA) on growth and development of these fungal strains considering possible fungicidal abilities of mentioned substances and comparing responses of fungi. The study shows inhibitory effect of ionones on mycelium growth of both Foln and Fo47. Our results also show the differences in apocarotenoid’s effect on studied strains in regards of sporulation, *FUB* genes cluster activity and fusaric acid (FA) production.

**Author summary:** In this study, we investigated the interaction between *Fusarium oxysporum*, a fungus that can either harm or potentially benefit plants, and natural plant-derived compounds known as apocarotenoids. We focused on two fungal strains: one that specifically infects flax plants, causing disease, and a nonpathogenic strain that may protect flax from pathogens. By examining the effects of apocarotenoids like ionones and abscisic acid, we aimed to understand how these compounds influence fungal growth, sporulation, toxin production, and gene activity related to pathogenicity. Our findings reveal that ionones inhibit the growth of both strains, suggesting their potential as antifungal agents. Interestingly, the two strains showed distinct responses to these compounds, particularly in their production of fusaric acid and activation of toxin-related genes. These results highlight the complexity of fungal interactions with plant-derived molecules and suggest that apocarotenoids could play a role in modulating fungal behavior. This work contributes to our understanding of plant-fungal interactions and may inform future strategies for managing crop diseases sustainably.

## Introduction

The fungi of the *Fusarium* genus are among the ’Top 10’ fungal pathogens that cause the greatest number of plant diseases [10]. One member of this genus, specifically the *Fusarium oxysporum* species complex, is among the most economically important. This complex consists of various forms that exhibit a high degree of specificity to their hosts. [24].

*Fusarium oxysporum* f. *lini* (Foln) is a soil-borne hemibiotroph that invades flax through the roots and causes a flax wilt [1, 2]. In a biotrophic phase, the fungus grows within root tip meristematic cells, causing cell death, cortex degradation, and hollowing of the root. Fungal growth eventually reaches protoxylem vessels, triggering the necrotrophic phase [17, 34]. Apart from a large number of pathogenic forms, non-pathogenic strains have also been isolated from the soil [24]. The nonpathogenic strain Fo47, a well-studied endophyte that can protect the plant from the pathogenic infections [15, 16]. Endophytes colonize plants mostly without reaching the xylem vessels, but remain in the area of the root cortex and endoderm [27]. *F. oxysporum* endophytes have been thought to induce endophyte-mediated resistance, which helps the plant from root invasion of pathogens [25, 26].

Strains from the FOSC, including non-pathogenic strains, can synthesize various mycotoxins, including fusaric acid (FA), fusarins, beauvericin, and moniliformin [3, 4]. Mycotoxins are dangerous primarily to animals, due to their carcinogenic and mutagenic effects [5]. For fungi, they are helpful in invading plants, especially fusaric acid [12, 18, 19]. FA is a natural alkaloid, specifically a 5-butylpicolinic acid, derived from aspartate (or oxaloacetate) and three acetate units [9, 11]. The biological effects of FA are concentration dependent. Toxic concentrations for plants exceed 10^-5^ M, leading to disruption of cell membrane potential reduced mitochondrial activity (inhibition of ATP synthesis); inhibition of root growth; induction of programmed cell death.

At non-toxic concentrations (e.g. 10^-6^ M), FA acts as an elicitor [4, 11, 12, 28, 30]. In *Arabidopsis thaliana*, FA induces the synthesis of phytoalexins, increases ROS production, elevates cytosolic calcium levels, and modulates ion channel currents [20]. In watermelon seedlings, FA inhibits root cell dehydrogenase activity, decreases cell membrane potential, and enhances lipid peroxidase activity, leading to increased malondialdehyde levels and activity of enzymes such as phenylalanine ammonia-lyase, catalase, superoxide dismutase, peroxidase, β-1,3-glucanase, and chitinase [8].

At least 12 genes from the *FUB* cluster are responsible for the synthesis of FA in the FOSC: five genes (*FUB1*, *FUB4,* and *FUB6-8*) are thought to be essential and *FUB10*, *FUB12* have a regulatory function [13]. FA production is also regulated with the specific velvet complex, which contains velvet proteins: VeA, VelB and VelC, and the LaeA. Proteins from this complex are also connected with fungal development [11, 43]. Fusaric acid production is also regulated by nitrogen and iron source availability and pH conditions [11, 48]. Some natural compounds can affect FA production. For example, sinapic acid, natural phenolic compound, increase FA production by *Fusarium oxysporum* f.sp. *niveum* [51].

Traditional agriculture methods primarily rely on chemical fungicides, which, while effective, pose several challenges, including the development of resistant fungal strains, environmental pollution, and potential health risks to humans and animals. The search for alternative solutions has led to increased interest in natural compounds with antifungal properties. These compounds, often derived from plants, bacteria, or other natural sources, offer a promising avenue for sustainable and eco-friendly fungal control [22].

This study aims to evaluate the antifungal effects of apocarotenoids against pathogenic strain *Foln* and nonpathogenic Fo47. By investigating the potency and spectrum of these compounds, we seek to identify potential candidates for development into natural fungicides.

Apocarotenoids are a group of compounds with various structures and functions. Some of them, such as ionones, are volatile. They are formed as a result of the oxidative decomposition of carotenoids catalyzed by carotenoid dioxygenases or by a non-enzymatic route. Among the apocarotenoids there are compounds with confirmed hormonal function (abscisic acid, strigolactones) and compounds for which signaling significance is only postulated, e.g. β-cyclocitral. Some earlier studies showed that these compounds may be involved in the response to environmental stress [14]. There is also a lot of research on the anticancer effects of ionones. The studies also showed chemopreventing, anty-inflammatory, cancer promoting and melanogenesis activity of these compounds [6]. Also, there are reports about antimicrobial and antifungal activity of apocarotenoids. For example, apocarotenoid β-ionone can inhibit the growth of microorganisms, including *Aspergillus flavus*, *A. parasiticus*, *A. fumigatus* and reduce the production of aflatoxins by *A. parasiticus* [6, 21, 32]. But there is a lack of information about the effect of apocarotenoids on fungal plant pathogens, such as *Fusarium oxysporum* or *Fusarium* as a genus at all.

## Results

### 1. Comparison of Characteristics Between Two Untreated Strains

The pathogenic and non-pathogenic strains show statistically significant differences in microspores and FA production as shown in **Table 1**. Fo47 produces 3-fold more microspores and practically 8-fold (in DW) and 30-fold more (in the medium) FA in comparison to Foln. Fo47 growth is 1,5-fold faster in comparison to Fol. . However the differences in fresh or dry weight of both strains was not observed.

**Table 1.**
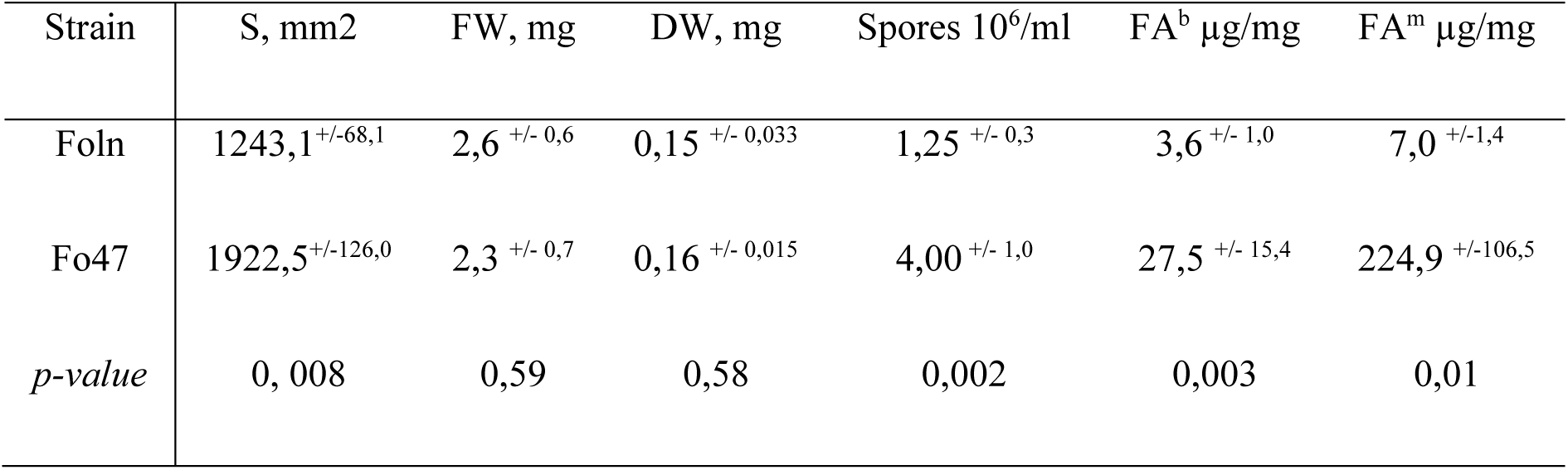
The comparison characteristics of Foln and Fo47 (FA^b^: FA content in fungal biomass, FA^m^ – in medium, µg/mg DW)

| Strain | S, mm<sup>2</sup> | FW, mg | DW, mg | Spores 10<sup>6</sup>/ml | FA<sup>b</sup> $\mu\text{g}/\text{mg}$ | FA<sup>m</sup> $\mu\text{g}/\text{mg}$ |
| --- | --- | --- | --- | --- | --- | --- |
| Foln | 1243,1<sup>+/-68,1</sup> | 2,6<sup>+/- 0,6</sup> | 0,15<sup>+/- 0,033</sup> | 1,25<sup>+/- 0,3</sup> | 3,6<sup>+/- 1,0</sup> | 7,0<sup>+/-1,4</sup> |
| Fo47 | 1922,5<sup>+/-126,0</sup> | 2,3<sup>+/- 0,7</sup> | 0,16<sup>+/- 0,015</sup> | 4,00<sup>+/- 1,0</sup> | 27,5<sup>+/- 15,4</sup> | 224,9<sup>+/-106,5</sup> |
| *p-value* | 0,008 | 0,59 | 0,58 | 0,002 | 0,003 | 0,01 |

### 2. Effect of Apocarotenoids on the Growth and Development of Foln and Fo47

#### 2.1 Mycelium growth

Macroscopic observations showed an inhibitory effect of ionones on the mycelial growth of both strains at all concentrations studied (**Fig.1**), which is supported with the colony size measurements (**Fig. 2**). Also, the lack of specific Fusarium pigment production and air mycelium growth is visible, but only at the added volume of 5 or 10 µL (Fig. S4-S6). The inhibitory effect of ionones on Fol is about 2-fold, but on Fo47 it is 4-fold. These results are statistically significant in all studied concentrations, excluding only 0,8 µL of a-ionone on Fo47 at 7 dpt. The effect persists also after 7 dpi. However, such effects was not observed for ABA treatment or β-cyclocitral (Fig. S7-S10). After these observations β- cyclocitral was excluded from tests. the higher concentrations of studied ionones was also excluded because, in further tests to determine their impact on fusaric acid production, the inhibition was so strong that the amount of the tested compound was beyond the detection limit. Comparing the growth speed of studied strains showed that Fo47 growth faster than Foln. All photos that show the effect of studied compounds at another timepoints or concentrations are located in the supplementary.

**Fig. 1:**
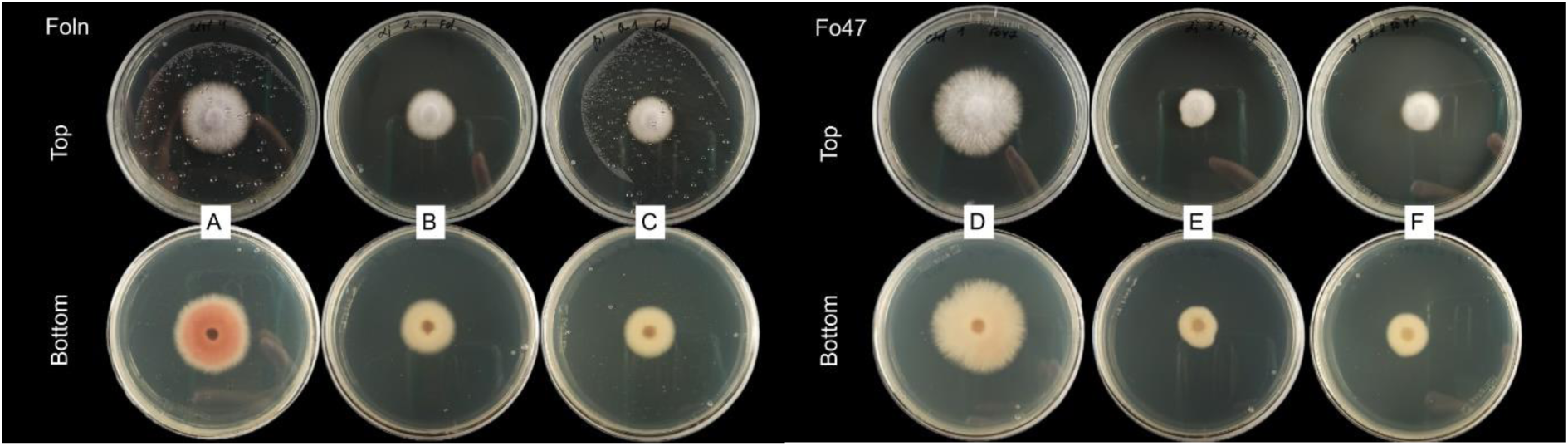
Effect of the ionones on the Foln (on the left side) and Fo47 (on the right side) mycelium growth; 3 dpt (A, D: untreated control; B, E: treatment with 2 µL α-ionone; C, F: treatment with 2 µL β-ionone). The images were taken from the top and the bottom of the Petri dish.

**Fig. 2:**
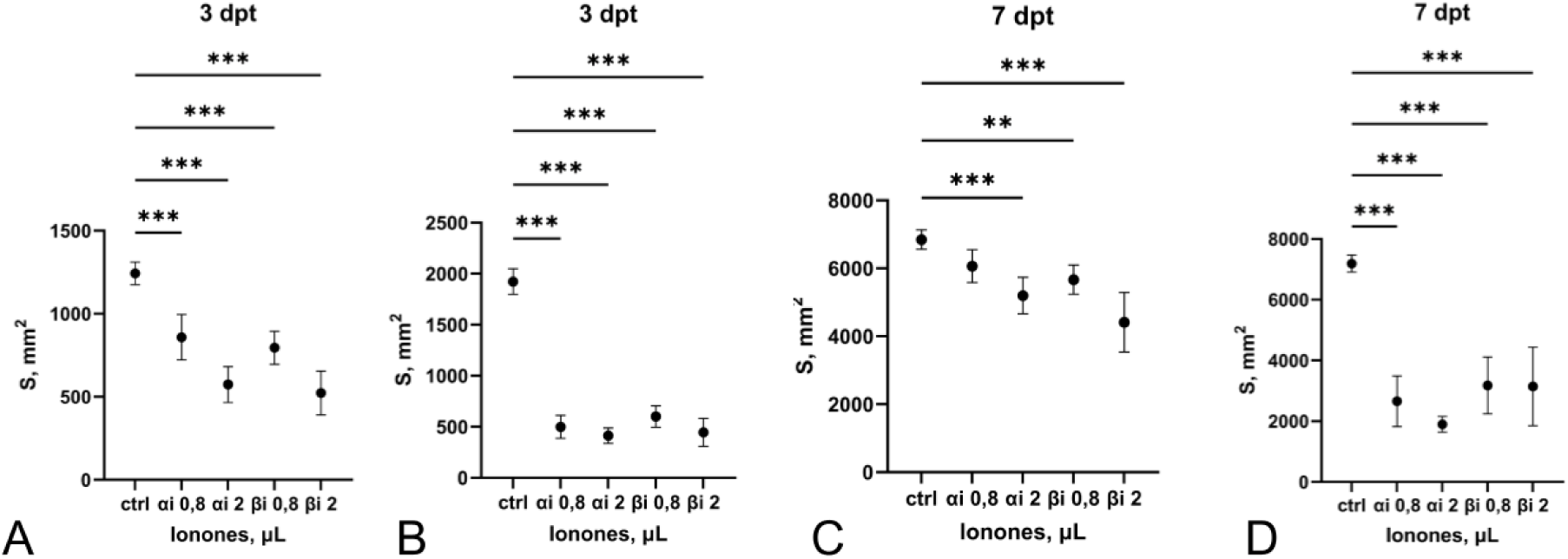
Effect of the ionones on the Foln (A, C) and Fo47 (B, D) colony size on 3d and 7th days after treatment with 0,8 and 2 µL of ionones.

#### 2.2 Microscopic observations

It was evident that ionones are increasing chlamydospores production in pathogenic strain, but not in Fo47. The accumulation of chlamydospores in treated cultures in comparison to single examples in nontreated ones was observed. Any effects of ABA treatment on the mycelium and spore structures were noticed.

#### 2.3 Microspores production

The stronger inhibitory effect of studied apocarotenoids on sporulation of the Fo47 in comparison to Foln was observed. Treatment with β-ionone decreased sporulation by about 2-fold. ABA treatment at 10 and 50 µL inhibited Fo47 sporulation by 1.5-fold compared to the untreated control. However, treatment with the lowest concentration of α-ionone (2 µL) showed significant 3-fold increase of the pathogen’s sporulation in comparison to untreated samples.

#### 2.4 Fungal biomass

There are no statistical differences in the fresh or dry weight of fungal biomass in liquid culture between nontreated and treated fungal strains (data in the supplementary(Fig.S2-S3)), so it was suggested that the tested compounds not affect the weight gain, but looking on the inhibitory effect on the mycelium growth it can be assumed that there are the difficulties in colony spreading and development.

##### 1. The level of mRNA transcripts of FUB and Velvet complex genes

The level of mRNA transcripts of FUB cluster genes and Velvet complex showed that the effect of studied compounds depends on concentration. The first heatmap (**Fig.5**) shows significant downregulation of Foln FUB genes (1-9) after α-ionone treatment, the cluster’s regulation genes are also downregulated. A similar effect observed after treatment with the higher amount of β-ionone, but the lower amount shows upregulation of a cluster. *LaeA* from velvet complex is upregulated after treatment with α-ionone, but v*eA* is downregulated, also after treatment with β-ionone.

**Fig. 3.**
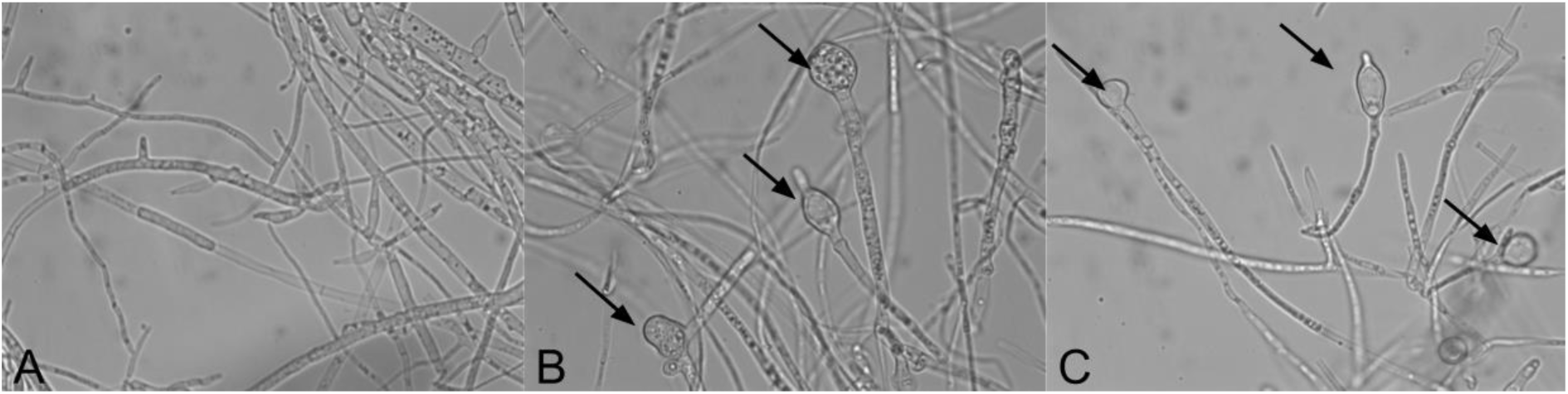
Chlamydospores formation in Foln: A - untreated control, B - treated with α-ionone; C - treated with β-ionone

**Fig 4.**
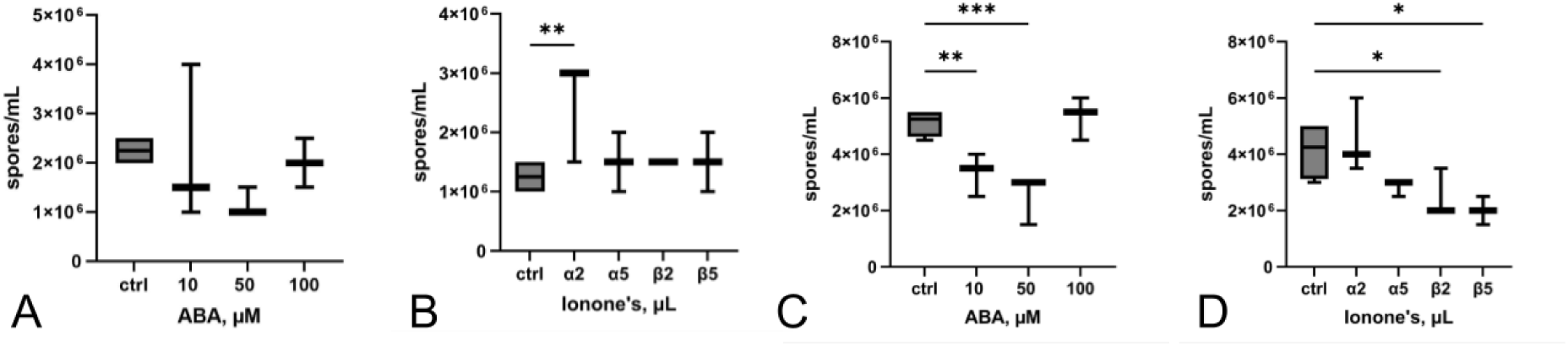
Effect of the ABA and ionones on the Foln (A, B) and Fo47 (C, D) microspore production; 3 dpt

**Fig 5.**
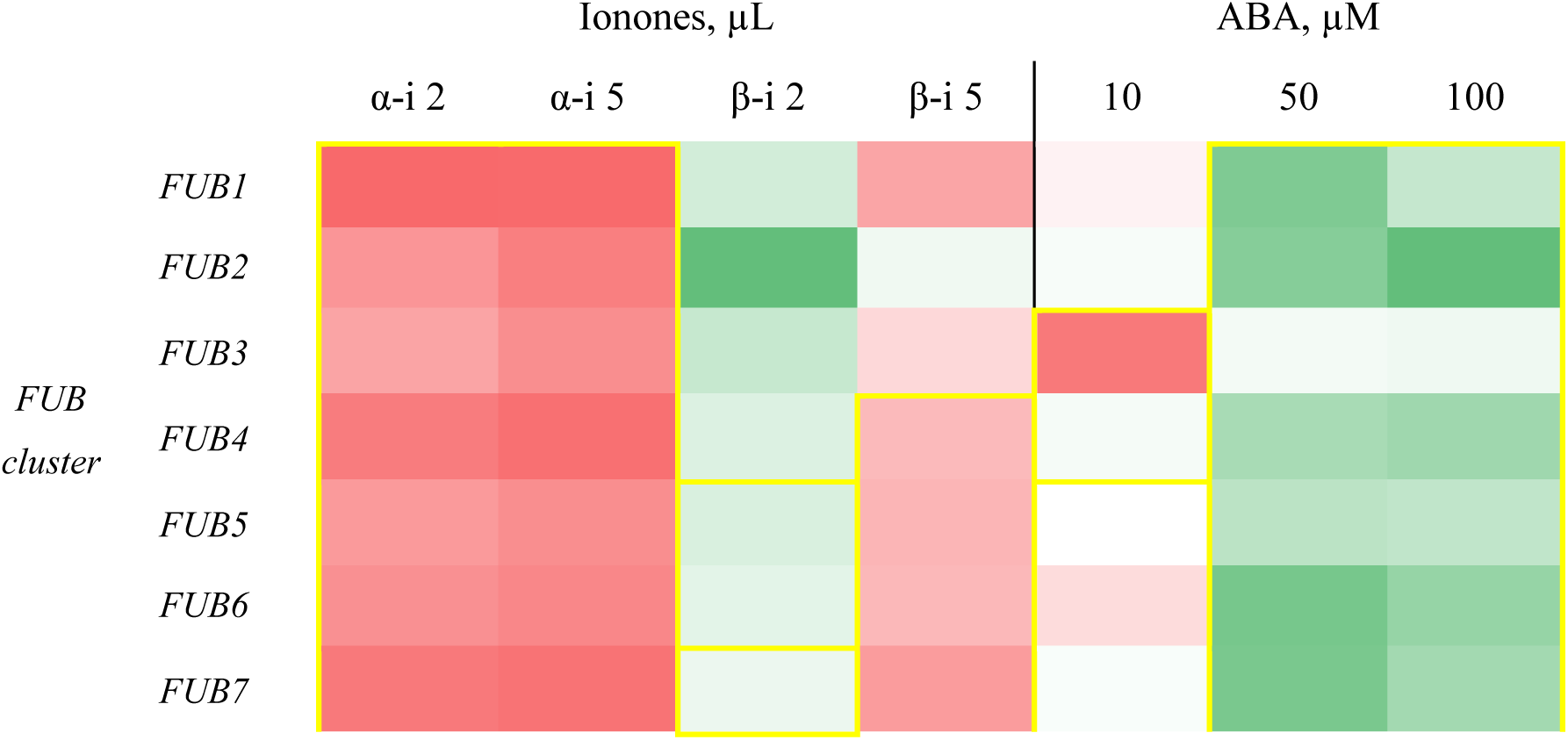
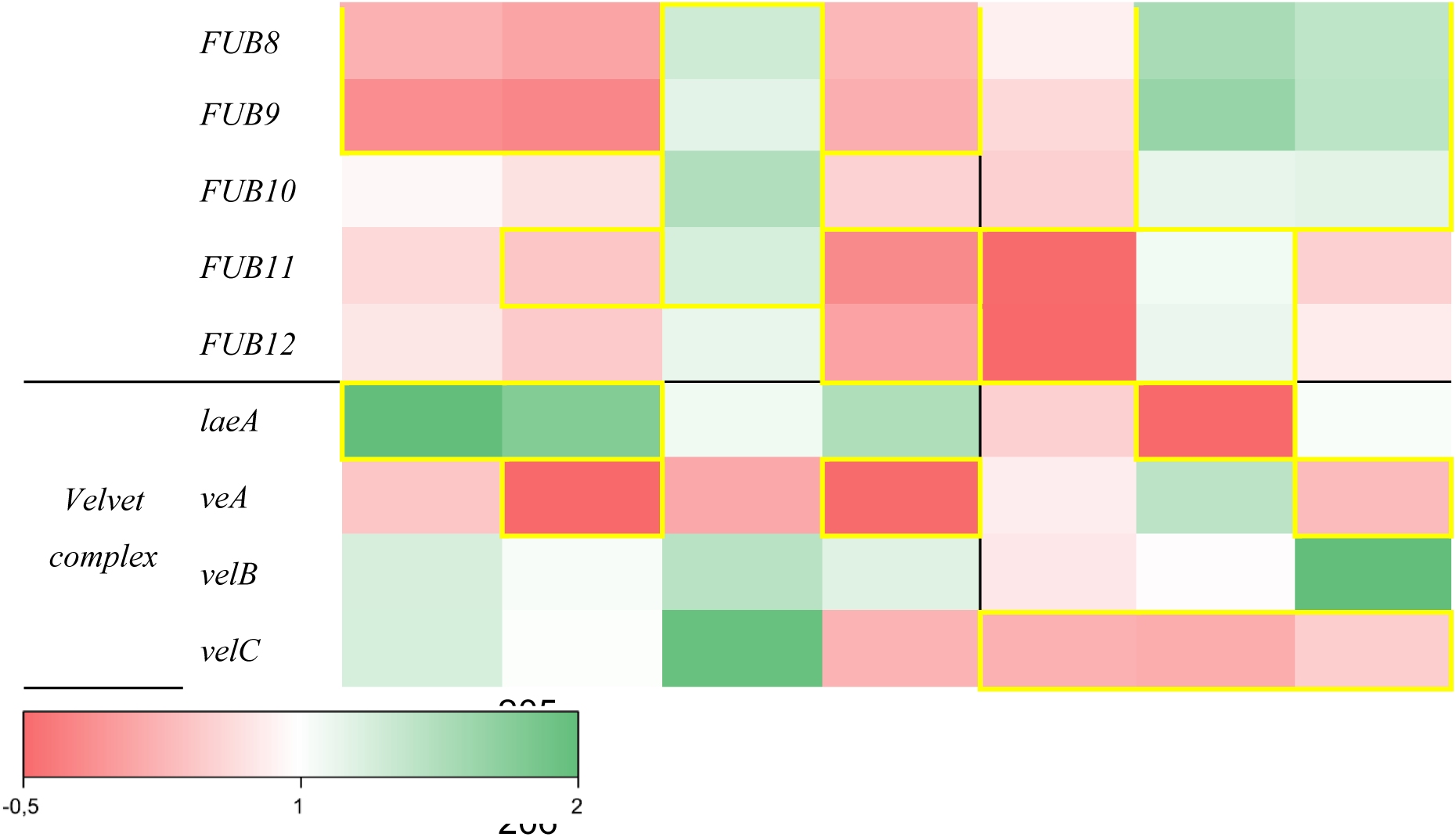
Heatmap of FUB cluster and velvet complex gene’s expression in Foln after ionones and ABA treatment; statistically significant data are framed in yellow. The exact data can be found in the supplementary (Table S3-S4).

The ABA effect is also concentration-dependent: all studied genes are striving to downregulation after 10 µM treatment, but higher concentrations activate FUB genes (1-9) and v*elB*. Interestingly, but only 50 µM ABA significantly downregulates *laeA.* The expression of *velC* decreased at every studied ABA concentration.

In comparison to Foln, the effect of studied compounds on the expression of Fo47 genes from FUB cluster and velvet complex is quite opposite (**Fig. 6**). Ionones upregulate all FA synthetic genes (*FUB1- 9*), but transporter (*FUB11*) and transcription factor (*FUB12*) are downregulated. *LaeA, veA* and v*elB* are also active, but v*elC* is downregulated. Taking into account regulation function of both *FUB12* and v*elC* and the results of FA production, it can be assumed that there is the downregulation of all genes at later time points.

**Fig. 6.**
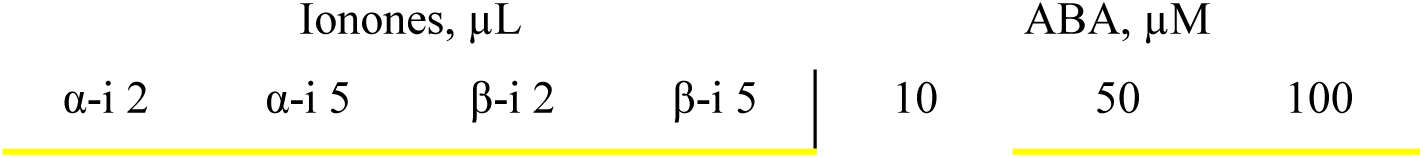
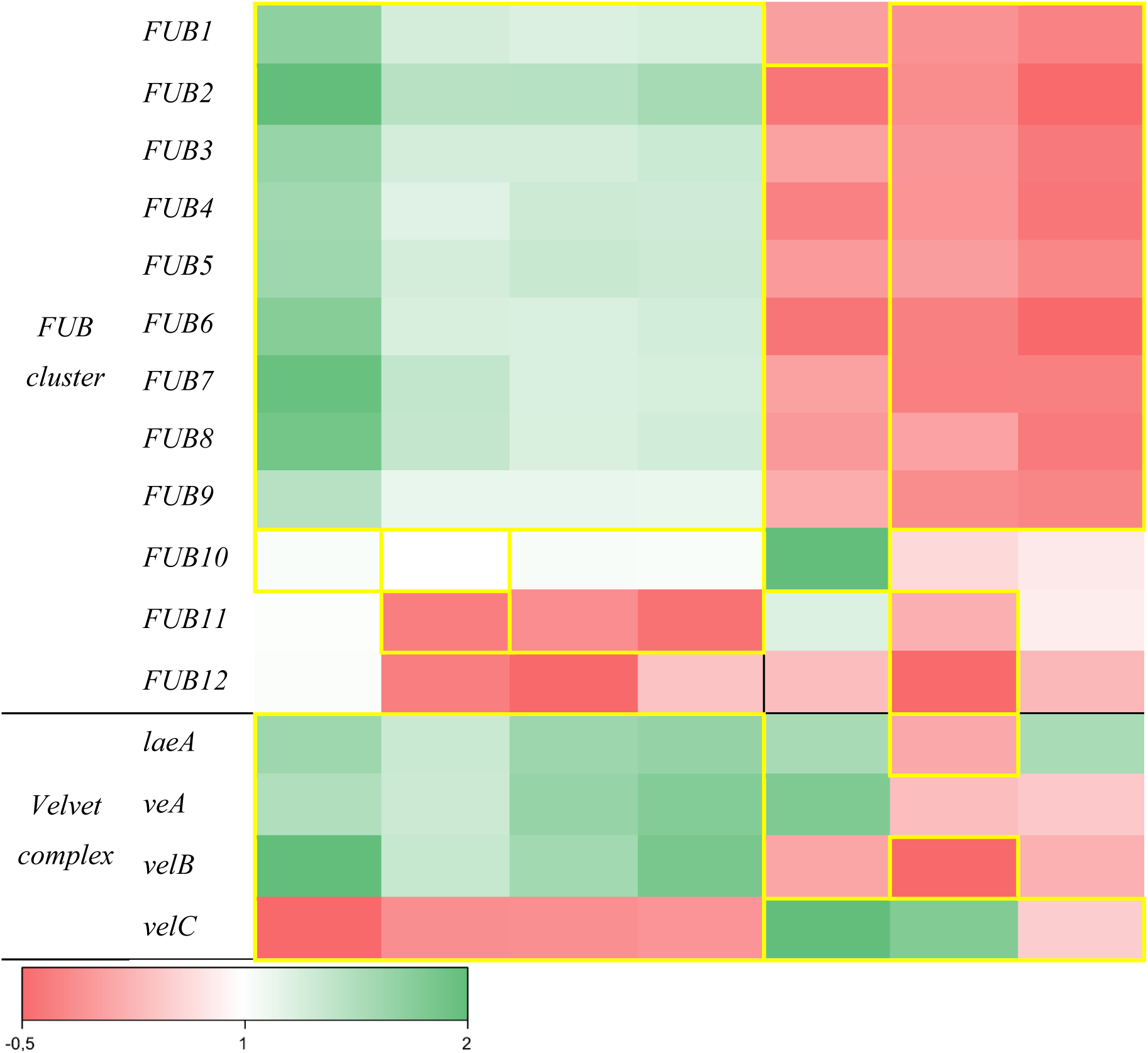
Heatmap of FUB cluster and Velvet complex gene’s expression in Fo47 after ionones and ABA treatment; statistically significant data are framed in yellow. The exact data can be found in the supplementary (Table S5-S6).

The ABA treatment downregulates activity of FUB cluster and velvet complex almost completely at that timepoint, but treatment with smaller concentration significantly activates regulation genes (*FUB10* and v*elC*).

##### 2. Fusaric acid production

The FA content in fungal biomass after 3 days of apocarotenoid’s treatment was checked. The gene expression analysis was also performed at this timepoint. And after 3 weeks it was checked whether the impact is sustained (in fungal biomass and in post-culture medium). It was found that the ionones start to have an effect on both the pathogen and nonpathogenic strain. These compounds have a strong inhibition effect on Foln FA production, that lasts for 3 weeks. Treatment with α-ionone resulted in a 5- to 10-fold reduction in FA production in biomass compared to untreated samples. In contrast, β-ionone exhibited lower efficacy, achieving only a 1.5- to 2-fold inhibition relative to the control. A similar tendency was observed in post- culture medium samples. Fo47 demonstrates low sensitivity to ionones, and this effect is not significantly reflected in the biomass results on 3^d^ week (**Fig. 7**). On the third day, a 1.5-fold inhibition of FA production was observed, and this effect remained consistent by the third week, showing a 1.5- to 2-fold inhibition FA production to the medium. In contrast treatment with ABA has and opposite effect. The ABA increases FA production in Foln from the beginning (1,5- fold) of treatment and lasts at least for 3 weeks (4- to 11-fold). Moreover, this increase practically reaches the level of FA production by non-treated Fo47 and is concentration dependent. However, ABA almost has no influence on FA production by Fo47 (**Fig. 8**).

**Fig. 7.**
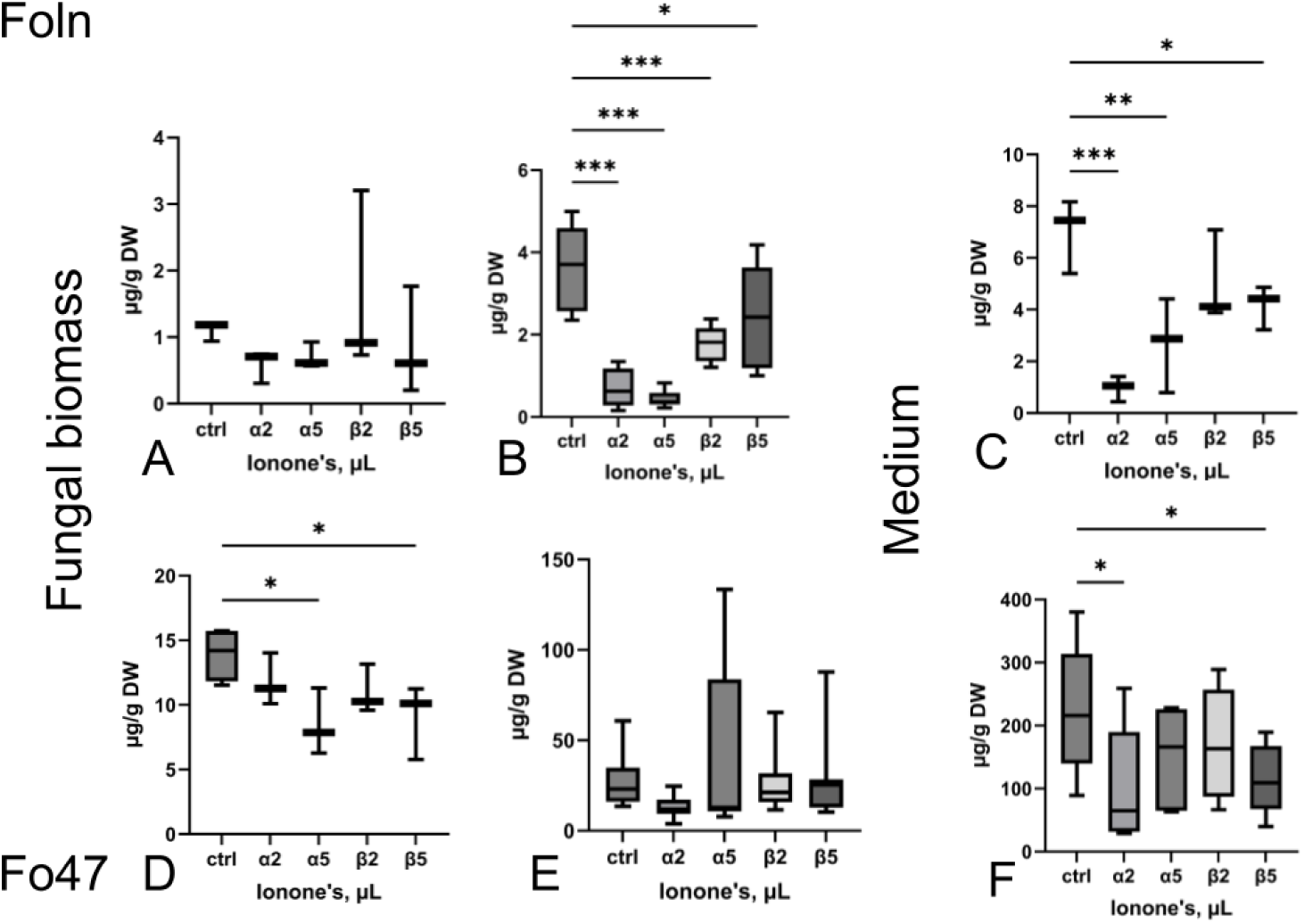
FA content in fungal biomass (A, D, - 3 dpt; B, E – 3 weeks) and in post-cultivation medium (C, F) after ionones treatment in Fol (A-C) and Fo47 (D-F)

**Fig. 8.**
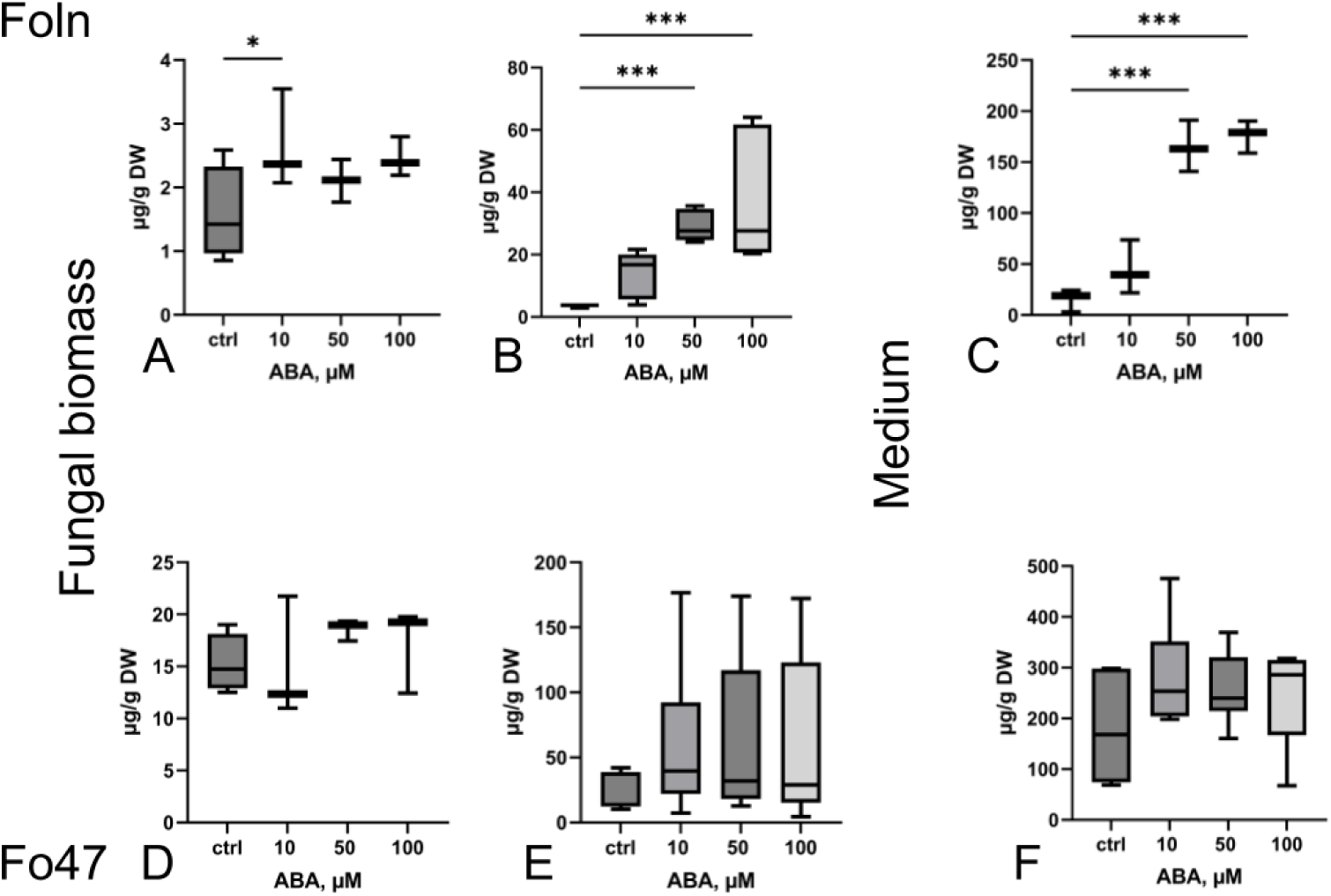
FA content in fungal biomass (A, D, - 3 dpt; B, E – 3 weeks) and in post-cultivation medium (C, F) after ABA treatment in Fol (A-C) and Fo47 (D-F)

## Discussion

The search for natural compounds to protect against fungal pathogens and molds is crucial in the context of increasing demand for organic and eco-friendly food production. A sustainable approach to preserving food safety and minimizing mycotoxin contamination is essential to meet consumer preferences. This highlights the importance of identifying natural compounds that can effectively inhibit fungal growth and reduce mycotoxin synthesis.

Natural compounds derived from the phenylpropanoid pathway, such as cinnamic acid, coumaric acid, and vanillic acid, have demonstrated diverse effects on fungal growth and mycotoxin production. These compounds have been shown to inhibit the growth of fungi, including *Fusarium oxysporum* f. sp. *lini* [7], *F. graminearum* [40, 41, 42], and *Monilinia fructicola*, as well as other phytopathogenic fungi like *Botrytis cinerea* and *Alternaria alternata* [24, 36]. Interestingly, phenolic compounds can have dual effects, either inhibiting or promoting mycotoxin production depending on the context. For example, treatment with cinnamic acid, coumaric acid, or vanillic acid increased the activity of hydrolytic enzymes, such as pectinase, cellulase, and amylase, in *F. oxysporum* f. sp. *niveum*, thereby enhancing mycotoxin production, including fusaric acid, while simultaneously inhibiting fungal growth and germination [24, 50, 51]. Certain alkaloids, such as capsaicin, and antifungal fractions from coffee bean extracts (which include flavonoids, carotenoids, and saponins) can effectively inhibit fungal growth and ochratoxin A production by *Aspergillus* species [37, 38]. This suggests that although phenylpropanoids may enhance fungal virulence factors, they also interfere with the fundamental processes required for fungal proliferation.

The effects of carotenoids and tocopherols on fungal growth and development remain relatively underexplored compared to other natural compounds. Notably, sub-lethal doses of α-tocopherol have been found to significantly reduce fumonisin production, while capsanthin has been demonstrated to inhibit aflatoxin production [23]. Picot et al. investigated the impact of several carotenoids on the growth and mycotoxin production of *Fusarium verticillioides*. The study shows the effects of lutein and zeaxanthin at a concentration of 0.7 μM, β-carotene at 0.4 μM, and α-tocopherol at concentrations ranging from 0 to 0.1 mM, finding no effect on fungal biomass. However, fumonisin production was affected, but only with α-tocopherol treatment [57]. This lack of effect on fungal biomass is consistent with our results. Boba et al. checked an influence of carotenoid extracts from transgenic flax with increased level of carotenoids on *F. oxysporum* f.sp *lini* and *F. culmorum*. The study indicated an inhibitory effect of these extracts on the mycelium growth of studied strains [58]. Interestingly, some studies have suggested that grains rich in carotenoids, such as oat grain, were found to be free of mycotoxins, while barley grains containing high carotenoid levels had a higher concentration of mycotoxins [31]. This paradox highlights the need for a more nuanced understanding of the relationship between carotenoids and fungal metabolism, as it appears that carotenoid-rich environments may not always offer protection against mycotoxin contamination. Cyclic terpenes, such as menthol, limonene, thymol at the concentrations of 25-1000 µL/L of culture medium inhibited the *F. verticillioides* growth and also fumonisin B1 synthesis [54, 55]. Essential oils, which contain diverse chemical compounds such as terpenes, alcohols, acids, esters, and aldehydes, have been shown to reduce the levels of mycotoxins like zearalenone and group B trichothecenes [22]. These findings indicate that certain natural compounds can modulate mycotoxin biosynthesis, offering potential avenues for controlling fungal contamination in food products.

The antifungal effects of ionones have been widely studied, particularly in terms of their impact on fungal growth and sporulation. Previous research by Wilson et al. demonstrated that β-ionone, applied in concentrations ranging from 10 to 1000 µL per liter of medium, inhibited both growth and sporulation of *Aspergillus parasiticus* and *A. flavus* [32]. Salt et al. observed that β-ionone suppressed the growth and sporulation of the tobacco pathogen *Peronospora hyoscyami* f.sp. *tabacina* [45]. Additionally, Felemban et al. showed that β-ionone inhibits the growth of *Botrytis cinerea* at millimolar concentrations but not at micromolar levels [47]. Synthetic derivatives of β-ionone have also exhibited interesting antifungal activity, with some promoting fungal growth (e.g., bromolactone derivatives promoting *F. oxysporum* f.sp. *lini* growth) while inhibiting others, such as *Aspergillus niger* [46]. Thiazolylhydrazone derivatives of β-ionone were found to exhibit antifungal activity against eight plant pathogens, including *F. oxysporum* f.sp. *niveum* and *F. verticillioides* [35]. Our results align with these findings: ionones significantly inhibited the growth of *F. oxysporum* f.sp. *lini* and reduced mycelial growth and sporulation in the nonpathogenic strain Fo47.

The differential response between pathogenic and nonpathogenic strains underscores the complexity of ionones’ effects and suggests that their antifungal mechanisms may vary based on the strain or species. Moreover, an accumulation of chlamydospores in response to ionone treatment was observed, which likely reflects unfavorable conditions for fungal growth. Chlamydospores play a crucial role in fungal survival, especially in colder environments, and are associated with the enhanced spread of flax wilt, as compared to microspores [44].

The velvet complex, which regulates both secondary metabolism and fungal development, including hyphal growth and sporulation, is also relevant to our findings. Deletion of key velvet complex genes, such as *veA*, *velB*, and *velC*, has been shown to reduce conidiation, aligning with our observations in the nonpathogenic strain Fo47. Similarly, the flat colony morphology and lack of aerial mycelium observed mutants ΔlaeA and ΔveA also correlate with our results, although this was only evident in Foln [43].

Regarding mycotoxin production, our study showed that ionones downregulated the expression of most genes from the FUB cluster in *F. oxysporum* f.sp. *lini*, resulting in reduced fusaric acid production. Interestingly, Foln and Fo47 exhibit nearly opposite responses of the tested genes at the observed time point. However, the nonpathogenic strain Fo47 was less susceptible to ionones in terms of fusaric acid biosynthesis, despite the inhibition of mycelial growth and sporulation. This suggests that ionones may target different pathways depending on the strain. Previous research by Wilson et al. also demonstrated the antifungal potential of β-ionone, showing its ability to reduce aflatoxin production [32].

The role of ABA in fungal growth and sporulation has been inconsistent across studies, depending on concentration and fungal species. Boba et al. reported that ABA at concentrations of 0.1 μM, 0.5 μM, 1 μM, 10 μM, and 50 μM had no observable effect on the growth or sporulation of *F. oxysporum* f.sp. *lini* and *F. culmorum* [7, 33]. Similarly, Buchrow et al. found that ABA, even at relatively high concentrations, had no effect on the growth of *F. graminearum* in both solid and liquid media. Additionally, ABA applications did not affect trichothecene gene cluster expression or deoxynivalenol production [49]. Petti et al. also demonstrated that ABA did not alter the growth of *F. culmorum*, regardless of the concentration [52]. Conversely, Qi et al. showed that ABA at concentrations of 475-950 µM inhibited mycelium growth of *F. graminearum* [53]. These discrepancies indicate that ABA exhibits varying effects on fungal growth and sporulation depending on the concentration and the specific fungal strain. Our findings support the notion that ABA has no effect on the growth or sporulation of *F. oxysporum* f.sp. *lini*, although the nonpathogenic strain showed inhibition of sporulation in response to ABA. The downregulation of the velvet complex after ABA treatment, however, did not correspond to changes in mycelium growth in our study, suggesting that the complex’s role may differ depending on the specific conditions or strains tested.

Regarding its effects on mycotoxin production, ABA appears to have strain-specific influences on secondary metabolism. Our study indicated a significant increase in fusaric acid production and FUB cluster expression in *F. oxysporum* f.sp. *lini* after ABA treatment. In contrast, the nonpathogenic strain exhibited downregulation of the FUB cluster, and fusaric acid levels remained unchanged. Xu et al. demonstrated that ABA affects both growth and secondary metabolite production in *A. nidulans* at 100 nM concentration [56]. These results suggest that the effect of ABA on mycotoxin production is also complex and may vary depending on the specific strain or species.

In conclusion, our research provides new insights into the effects of apocarotenoids, particularly ionones and ABA, on two strains of *Fusarium oxysporum*. Ionones were found to inhibit growth and reduce mycotoxin production, while ABA exhibited more variable effects, depending on concentration and fungal strain. These findings highlight the potential of ionones as biopesticides and suggest that further research is needed to explore their mechanisms of action in greater detail.

## Materials and Methods

### Biological material

The non-pathogenic strain *Fusarium oxysporum* 47 (ATCC Number: MYA-1198) and the pathogenic strain *Fusarium oxysporum* f.sp. *lini* (Bolley) Snyder et Hansen (ATCC MYA-1201) were purchased from ATCC (USA).

### Fungal Strains Cultivation and Experimental Conditions

Fungal strains were cultivated under varying conditions depending on the type of analysis performed.

- **Pre-culture preparation:** The fungal strains were grown for one week at 28°C in darkness on Petri dishes containing 20 mL of PDA medium and inoculated with an evenly distributed spore solution with the amount of 2×10^5^ spores.
- **Morphological Analysis:** A 3 mm plug from a one-week-old fungal preculture was placed in the center of the plate. The fungal strains were grown on Petri dishes containing 20 mL of PDA medium at 28°C in darkness. Morphological observations were conducted between 3- and 14- day post-treatment. The measurements of fungal colonies were performed using images and ImageJ software. The calculations of the colony size were performed using formula

S = π*ab* mm^2^

where, *a* is the length of the semi-major axis and *b* is the length of the semi-minor axis.

- **Spore production and the level of mRNA transcripts:** For these analyses, the strains were cultivated in 50 mL of CDB medium under the same conditions of 28°C in darkness, with shaking at 100 rpm for one week. A 3 mm plug from a one-week-old fungal preculture was inoculated into the medium. Apocarotenoids were added, and samples were collected after 3 days post-treatment. Liquid culture samples were separated into fungal tissues and post-cultivation medium and stored at -80°C. After that the samples were lyophilized and analyzed. Fusaric acid content in fungal biomass was also evaluated.
- **Spore’s content measurements**: The spores were measured in post-cultivation medium. Each sample was centrifuged at 3000 *g* for 10 min, the medium was discarded, and the spores were resuspended in 1 ml of distilled water. After that, 10 µL of suspension were placed on Thoma calculating chamber. Spores were calculated in 8 small squares.
- **FA Production and Mass Measurements:** This experiment was performed as in the case of defining spore production and genetic analysis, but with some changes: cultures were treated right away and were grown for 3 weeks, and FA content was also checked in the post-cultivation medium.

### Apocarotenoid Treatments

The antifungal activity of α-ionone, β-ionone, β-cyclocitral and abscisic acid (ABA) was tested. The volatile compounds were added to the culture medium in volumes of 0,8 µL and 2 µL for 20 ml of PDA medium, or 2 µL and 5 µL for 50 mL of CDB medium. It is an equivalent to concentrations ranging from 40 to 100 µL/L of medium (Table S2). Untreated cultures served as the control group. For ABA we tested concentrations of 10 µM, 50 µM and 100 µM, using the highest solvent concentration (methanol) as a control. All apocarotenoid compounds were purchased from Sigma-Aldrich.

### Microscopic observations

Microscopic observations were performed under the epi-fluorescent microscope (ZEISS AXIO Scope A.1.) at 40x magnification. Images were documented using the black and white camera AxioCam ICm1 and ZEISS ZEN 2.6. lite software.

### In *silico* analysis and Primer’s design

In order to find out which FA synthesis and regulatory genes from the *FUB* cluster and Velvet complex are present in the strains we studied, the previously published genome of *Foln* (txid120040) and Fo47 (txid660027) was analyzed and, based on homology the sequences of interest were selected. Based on the *in-silico* analysis and the NCBI primer Blast tool, appropriate primers were designed (Table S1).

### House-keeping gene selection

In order to select the appropriate reference gene, qPCR was performed on a BioTad thermal cycler, and the results were analyzed using CFX Maestro Software for CFX Real-Time PCR Instruments (BioRad). For the analysis three potential house-keeping genes were tested: ubiquitin (*UBI*), β-tubulin (*TUB2*) and elongation factor 1α (*EF1A*) (Fig. S1).

### The level of mRNA transcripts

The level of mRNA transcripts was measured to determine expression of the *FUB* genes cluster and Velvet complex which are involved in the production of the FA were assessed by the Real Time PCR method. Total RNA was isolated using the GeneMATRIX DNA/RNA Extracol Kit (EURx) according to the attached protocol. The quality of the isolated RNA was examined by means of electrophoresis in denaturing conditions. The quantity and quality of the isolated RNA were checked using Implen NanoPhotometer. Isolated RNA, after removing DNA residues (with DNase I (ThermoScientifc) as described in the manufacturer’s protocol), were serve as the template for cDNA synthesis using a High-Capacity cDNA Reverse Transcription Kit (Applied Biosystems). PCR reactions in real time are carried out using SG qPCR Master Mix (EURx) in a thermocycler StepOnePlus™ Real-Time PCR system from Applied Biosystems. Changes in gene expression levels were presented as relative amounts with respect to a reference gene (*UBI*). The reaction conditions were: 50 °C for 2 min, 95 °C for 10 min (holding stage); and 95 °C for 15 s, 57 °C for 30 s, 72 °C for 30 s, 40 cycles (cycling stage). The conditions of the melting curve stage were: 95 °C for 15 s, 60 °C for 1 min, 95 °C for 30 s, and ramp rate: 1.5%. Real-time PCR reactions were performed in triplicate for each of the analyzed samples.

### Determination of the FA content

20 mg of the lyophilized fungal tissues were used for the FA extraction. 4 ml methanol were added to the samples with 100 µL of salicylic acid methanol solution (1 mg/ml) as internal standard (IS). The samples were placed in an ultrasonic bath for 1 hour and then placed into shaker for 4 hours in 12 °C. After filtration with PTFE Acrodisc Syringe Filter, 0.45 μm 25 mm (Waters), extracts were dried using SpeedVac^TM^ Refrigerated Centrifugal Vacuum Concentrator (ThermoScientific^TM^) and resuspended in 1 ml UPLC grade methanol. Extracts were filtered using PTFE Acrodisc Mini Syringe Filter, 0.2 μm 13 mm (Waters). The extracts were analyzed with ultra-high performance liquid chromatography (UPLC) with a diode array detector (Waters Acquity UPLC system). The separation was carried out on the ACQUITY UPLC BEH C18 Column, 130Å, 1.7 µm, 2.1 mm X 100 mm. We used the method described by *Boba A. et al* with no changes, for the compound’s separation. The analyzed compounds were identified on the basis of retention -fold and spectra, and then the amount was determined on the basis of adequate standard chromatograms [7].

### Statistical analysis

Statistical analysis was performed using GraphPad Prism 10 and Statistica 13 software (StatSoft, USA). Oneway analysis of variance ANOVA with Fisher’s posthoc test was used. For the Foln and Fo47 comparison t-test was used. Statistically significant differences are marked: * for p < 0.05, ** for p < 0.01 and *** for p < 0.001.

## Authors contributions

YK- Investigation, Methodology, Writing – Original Draft Preparation, Visualization, Formal Analysis, Writing – Review & Editing

MBS- Formal Analysis

AB - Funding Acquisition, Project Administration, Supervision, Methodology, Writing – Review & Editing, Resources

MP- Resources

JMD- Funding Acquisition, Supervision, Methodology, Writing – Review & Editing, Resources

AK – conceptualization, Funding Acquisition, Project Administration, Supervision, Methodology, Re- sources , Writing – Review & Editing

## Acknowledgments

The work was in part financed by National Science Centre (NCN), Grants Nos. NCN 2018/29/B/NZ9/00288, 2022/47/B/NZ9/00998 and 2022/47/D/NZ9/00642.

## Abbreviations

Foln: Fusarium oxysporum f. sp. lini
Fo47: Fusarium oxysporum strain 47
FOSC: Fusarium oxysporum species complex
FA: fusaric acid
ABA: abscisic acid α-i - α-ionone
β-i: β-ionone
Fig.: Figure
PDA: potato dextrose agar
CDB: Czapek-Dox Borth
FW: fresh weight
DW: dry weight
Hpt: hours past treatment
Dpt: days past treatment

